# Pumping stations negatively affect the distribution of critically endangered European eel (*Anguilla anguilla*); a landscape-scale study using environmental DNA metabarcoding

**DOI:** 10.64898/2026.08.28.746846

**Authors:** Angus I.T. Monaghan, Nathan P. Griffiths, Graham S. Sellers, Lori Lawson Handley, Andy D. Nunn, Bernd Hänfling, James A. Macarthur, Rosalind M. Wright, Marco Cattaneo, Jonathan D. Bolland

## Abstract

**Context:** Pumping stations pose a threat to fish globally through land use change, habitat fragmentation and entrainment risk, with the catadromous and critically endangered European eel particularly impacted.

**Objectives/methods:** Establish, model, assess and understand the present-day distribution of European eel and resident fishes in 152 pumping station catchments in a once extensive wetland (The Fens) using eDNA metabarcoding (855 samples over two and half years), with specific focus on anthropogenic influences on hydrological connectivity and habitat quality. A removal survey design maximised confidence in negative results while minimising time and consumable costs.

**Results:** Eel occurrence upstream of pumping stations was low (occupancy = 28.3%) and positively associated with catchment area, fish species richness and natural hydrological connectivity (gravity drainage or flooding) and negatively associated with distance from the tidal limit. Fish species richness replaced catchment area and improved model performance, potentially acting as a biotic indicator of habitat quality and connectivity. Pumped catchments with manually operated upstream water transfers had reduced eel presence, potentially linked to the direction of water flow or the timing of operation. By contrast, fish species richness increased in these catchments during summer, suggesting displacement into unsuitable long-term habitats. Physical habitat maintenance had no detectable effect on eel occurrence or fish species richness.

**Conclusions:** This study provides the first landscape-scale assessment of European eel distribution and drivers of occurrence in pumped river catchments. The highly novel and comprehensive insights have implications for European eel conservation as well as infrastructure and catchment management, including compliance with legislation (EC Regulation No. 1100/2007).

## Introduction

The European eel (*Anguilla anguilla*) is a catadromous fish species with a native freshwater range that includes much of Europe, certain Mediterranean/Atlantic areas of North Africa and Mediterranean Western Asia (Pike et al., 2020). Listed as ‘Critically Endangered’ on the International Union for the Conservation of Nature (IUCN) Red List of Threatened Species, it is a species of high conservation priority given an estimated 95% decline in recruitment in the last 50 years (Pike et al., 2020). Indeed, the decline is thought to have been due to a number of factors including: climate change (Bonhommeau et al., 2008), overfishing (Baisez & Laffaille, 2008), habitat fragmentation (Dekker, 2000; Podda et al., 2022), migratory barriers (Piper et al., 2013, 2017), pollution (Corsi et al., 2005) and mortality induced by entrainment in water-control structures (Bolland et al., 2019). Possessing a complex, varied life-cycle, the European eel has very particular, often challenging, requirements for its effective conservation (Jacoby et al., 2015; Verhelst et al., 2021; Mukherjee et al., 2026). To aid their protection, the EU introduced legislation for conservation measures with the development of Eel Management Plans (EC Regulation No. 1100/2007; European Commission, 2007). The Eels (England and Wales) Regulations 2009 has specific legislation that requires any intakes abstracting >20 m^3^ of water per day, including pumping stations, to install European eel protection measures (unless exempted by the Environment Agency).

Pumping stations are essential infrastructure for flood defence in low-lying areas and reclaimed land but represent a range of ecological threats, especially to fish. Downstream passage at pumping stations is a significant issue for mature, silver eels attempting to leave fresh water to return to the Sargasso Sea to spawn. Injury and mortality during entrainment through pumps are a particular concern (Buysse et al., 2014; Bolland et al., 2019; van Keeken et al., 2021), as well as migratory delays and disruptions to silver eels that do manage to navigate pumping stations (Bolland et al., 2019; Evans et al., 2024a, 2024b). Resident fish are also at risk, especially during pump start-up and extreme flood-relief operation, when flows frequently exceed fish swimming capabilities (Norman et al., 2023a, 2023b, 2024). Notwithstanding, pumping stations and associated infrastructure (e.g. flood banks/dikes and sluices) are also barriers to upstream passage (Rolls et al., 2014; Deflem et al., 2022), and may totally exclude eels from entire catchments they inhabited historically (Griffiths et al., 2025a). Some pumped catchments, however, have periodic improvements in hydrological connectivity via gravity discharges, over-bank flooding or manual water transfers (e.g. to maintain river levels for agricultural irrigation), which have implications for the prevailing fish community.

River maintenance measures, which are typically both extensive and intensive in pumped catchments, also present many threats to eels and wider fish communities. Habitat complexity is reduced and watercourses are often transformed into uniform drainage channels (Van Loon et al., 2009) with no flow, except when pumps are operating, and thus lentic fish communities prevail (Townsend & Peirson, 1988; Buisson et al., 2008). As such, the presence of rheophilic or marine fish species in pumping stations that discharge into freshwater and tidal water, respectively, can be used as evidence of connectivity between a pumped catchment and its receiving waterbody. For example, regular transfers for irrigation can lead to specialist fish species being relocated into unsuitable habitats (Roberts & Rahel, 2008). Furthermore, drainage channels are often prone to seasonal drought (Herzon & Helenius, 2008) and eutrophication (Janse, 2005) and, as a result, dissolved oxygen sags, which can be extremely harmful to fish (Dawson, 2002). Without long-term or high-frequency water quality monitoring, as is the case for pumping stations studied here, episodic water quality events may go undetected or be misinterpreted. Following fish kills, recolonisation from downstream may be prevented, and thus impoverished fish communities may persist. Consequently, the prevailing fish community can act as long-term biotic indicators of ecosystem health (Blabolil et al., 2017) and connectivity (Sun et al., 2022) given the prevalence of physical habitat maintenance measures and highly variable hydrological connectivity in pumped catchments.

In fragmented systems, eels are often rare (low site occupancy) and present in low abundance (Griffiths et al., 2020), and establishing distributions confidently is therefore challenging (Griffiths et al., 2025b). Recent advances have occurred in the field of environmental DNA (eDNA) metabarcoding, i.e., the analysis of traces of genetic material in environmental samples, and it is recognised as an effective method of monitoring aquatic and terrestrial communities, often out-performing traditional sampling methods (Hänfling et al., 2016; Weldon et al., 2020; Penaluna et al., 2021). Indeed, eDNA metabarcoding has proven effective for monitoring the distribution of eels in pumped catchments (Griffiths et al., 2020). It is also possible to assign confidence in absence when a target species is not detected, and thus mitigate the longstanding uncertainty around false negative results when using eDNA-based monitoring (Griffiths et al., 2025b).

To effectively conserve European eels, there is an urgent need to establish and understand their present-day distribution, especially in understudied habitats that have been subject to considerable land-use change. In this study, a landscape-scale assessment of European eel and resident fish distributions upstream of pumping stations in a once extensive wetland was performed using eDNA metabarcoding. The confidence in European eel absence was quantified, and their occurrence was modelled and assessed against abiotic (distance from tidal limit, catchment area, hydrological connectivity and habitat management) and biotic (resident fish community) variables. Fish species richness was also modelled and assessed against the same abiotic variables, given their potential to act as biotic indicators of habitat and water quality and connectivity. Throughout, particular attention was given to the type and timing of hydrological connectivity, including the incorporation of rheophilic and marine species as proxies for upstream connection, as well as the type and frequency of habitat management. These highly novel and comprehensive insights will have implications for European eel and resident fish conservation as well as infrastructure and catchment management in anthropogenically dominated landscapes.

## Materials and methods

### Survey design

This study was carried out on pumped catchments in the River Great Ouse and River Nene basins, England, part of what is locally known, and is referred to hereafter, as The Fens. Once an expansive wetland, The Fens was drained in the 17th century and now comprises approximately 4000 km^2^ (Smith et al., 2010) of mostly agricultural land with a complex network of heavily modified drainage channels controlled by pumping stations (Solomon & Wright, 2012). Here, 152 pumping station catchments in The Fens were studied using a removal survey design described by Mackenzie & Royle (2005), where multiple rounds of sampling took place but only sites negative for the target species (European eel) were re-sampled, thereby maximising confidence in negative results while minimising time and consumable costs. Notwithstanding, there were five instances when samples with <50 eel sequencing reads were re-sampled in the following round to provide additional confidence in the assessment. Water was sampled in four rounds; a single water sample was collected in May 2021 (R1) and two water samples were collected in March 2022 (R2), July/August 2022 (R3) and September 2023 (R4). However, there were exceptions and revisions due to the large-scale, long-term nature of the study. Specifically, seven “high priority” pumping stations had ten samples collected in R1 and were not resampled thereafter due to the urgent need for a highly confident eel presence/absence verdict prior to imminent refurbishment (Figure 1). Twenty-eight sites with very poor habitat quality (i.e. river width, depth and length) and a “poor” (<4) fish species richness in R1 were deemed unsuitable for eel colonisation, and thus were not sampled in R2. Nine of these sites were reincorporated from R3 (Figure 1) and five had four samples taken in R4 to ensure no potential eel sites were missed. Twenty-six eel-negative sites that were inhibited in R1 also had an additional sample taken in R4 (Figure 1).

**Figure 1.**
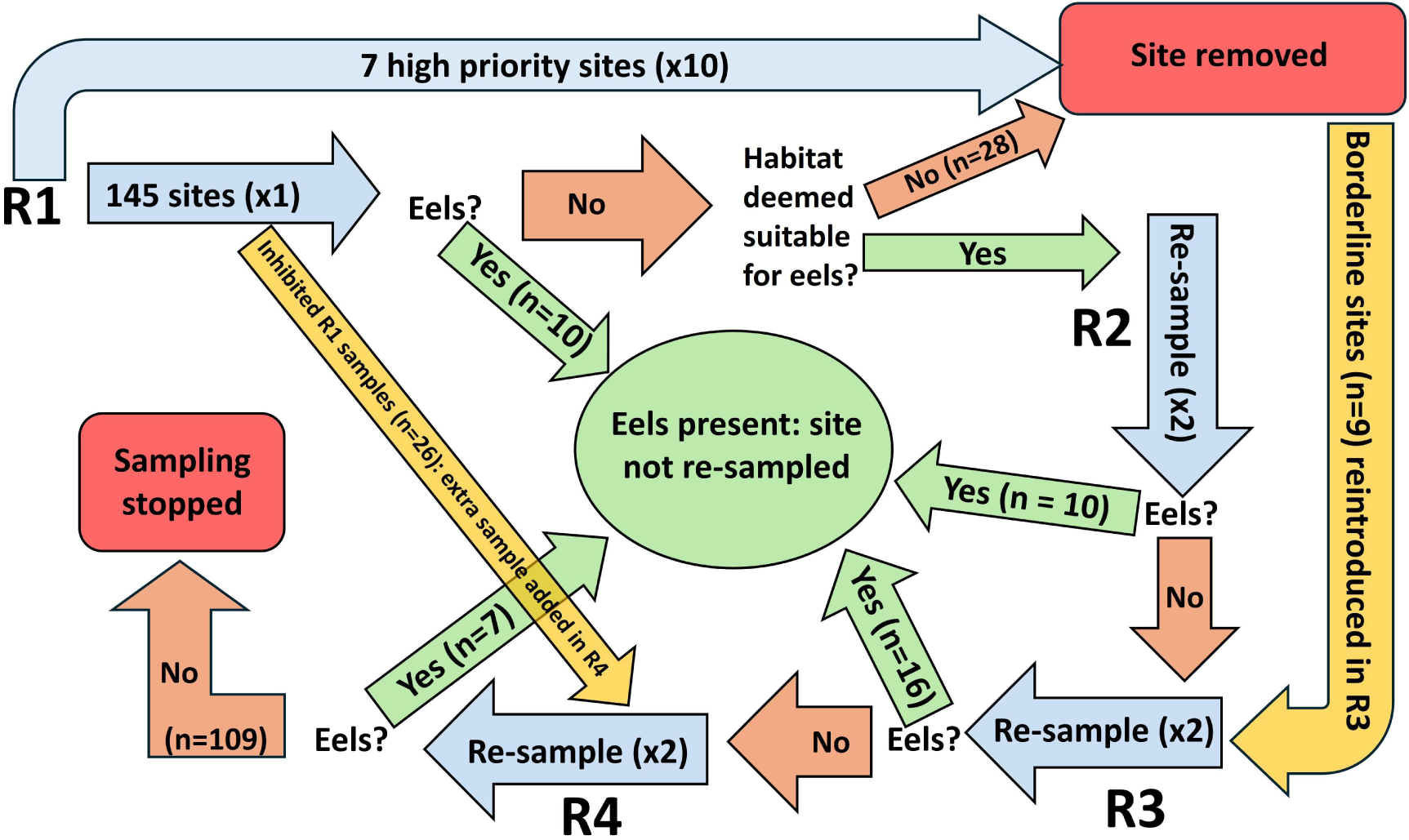
Flow chart detailing the sampling design and rationale.

### Sample collection and filtration

Two-litre surface-water samples were collected in sterile bottles (Gosselin HDPE plastic bottles, Fisher Scientific, UK) directly upstream of each pumping station following protocols outlined in Griffiths et al. (2025a). The rationale for sampling at the structures, aside from consistently reliable access, was that it is typically the widest and deepest part of the catchment and aquatic eDNA will be transported downstream and accumulate near pumping stations, and thus provides the best single sampling point to represent the species present upstream. Three sub-samples of approximately equal volume were taken per bottle to account for potential heterogeneity of eDNA distribution in the water column. All equipment was sterilised with a 10% sodium hypochlorite bleach solution and rinsed thoroughly before each use. Field blanks (1 per day) were employed throughout sampling. After collection, all samples were placed on ice until arrival at the laboratory, where the bottles were sterilised with 10% bleach solution and stored in a clean refrigerator at 4 ℃ overnight.

All samples were filtered through sterile 0.45 μm cellulose membrane filters (Whatman, UK) using vacuum pumps within 24 hrs of collection. Two filters were used per sample to minimise clogging. Filters were changed either when the first litre of the sample had been filtered or after 30 minutes – whichever happened first. After filtration, the filters were stored in sterile Axygen 5 ml screw-cap tubes at −20 °C until needed for DNA extraction. Between each sample, filtration equipment was sterilised with 10% bleach solution for 10 minutes, followed by 5% lipsol for 5 minutes (to remove bleach residue), and then rinsed with purified water.

### DNA extraction

DNA was extracted according to the Mu-DNA water extraction protocol described by Sellers et al. (2018) and, following R1, with the addition of an extra ethanol wash step to minimise the risk of PCR inhibition. Extraction blanks (1 per day) were employed.

### DNA metabarcoding

DNA metabarcoding followed the two-step nested PCR protocol described in Griffiths et al. (2023), using vertebrate-specific 12S-V5 primers (Riaz et al., 2011) to amplify a 106 bp fragment under the cycling conditions described therein. Libraries were sequenced on an Illumina MiSeq and sequences were demultiplexed using a custom bioinformatics workflow (https://github.com/EvoHull/Tapirs) and assigned against a curated UK vertebrate database using a 98% identity threshold. The 12S assay does not distinguish between perch (*Perca fluviatilis*) and zander (*Sander lucioperca*), therefore all detections were reported as Percidae. Detailed descriptions of DNA metabarcoding workflows for these samples can be found in Monaghan et al. (2026).

### Data preparation and analysis

#### Covariate preparation

Catchment areas of pumping stations and their distances from the tidal limit was taken from Solomon & Wright (2012). Catchment areas were natural log-transformed for analyses due to the data spanning several orders of magnitude. The number of barriers between pumping stations and tidal water were identified from satellite imagery. Hydrological connectivity and habitat management information was collected for all but three sites via contact with pumping station operators; analyses using these data were restricted to 149 sites.

Each pumping station catchment was categorised according to the type of hydrological connectivity and thus the viability for upstream European eel passage into the pumped catchment from a downstream waterbody or neighbouring catchment (Table 1; Figure S1). Higher-numbered connectivity types were hypothesised to be better connected for upstream migrating eels. Sites with multiple connectivity types were assigned the highest number. These five connectivity types were also combined into three connectivity groups, i.e. none, manual and natural, with none and manual also grouped into non-natural. Type 0 sites (n = 24) had no known connectivity to other catchments other than downstream movement of water through pumps during operation. Type 1 (n = 3) and Type 2 (n = 96) represent pumped and non-pumped transfers of water into a pumped catchment, respectively, which are grouped into manual connectivity. Manual connectivity (Types 1 and 2) primarily occurs to replenish water levels in pumped catchments used to irrigate arable fenland during drier periods, i.e. mid-to-late summer and early autumn. Sites with manual connectivity were also categorised according to whether water was transferred from the downstream waterbody (n = 82) or neighbouring catchment (n = 17). There was too little variation in the frequency of manual water transfers to elucidate any influence; all but three sites that transferred water did so multiple times per year (Table S1). Type 3 (n = 9) sites were connected during floods from downstream/neighbouring catchments and Type 4 (n = 17) sites had gravity drainage from the pumped catchment, which are grouped into natural connectivity. During comparisons, none and manual are also grouped into non-natural.

**Table 1.**
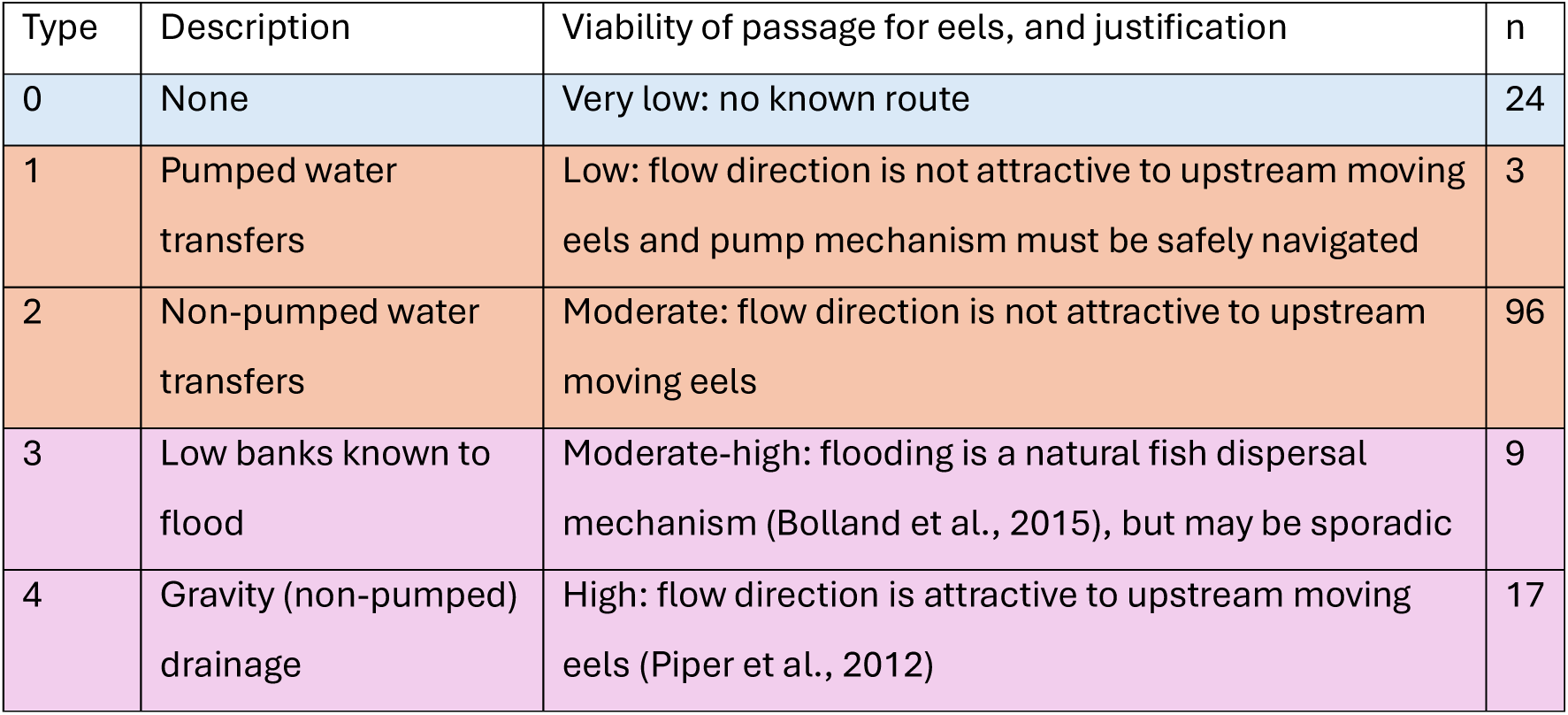
Hydrological connectivity types according to the viability for upstream European eel passage into the pumped catchment from a downstream waterbody or neighbouring catchment; higher scores represent better connectivity for eel. Types are also grouped according to whether there is none (blue), manual (orange) and natural (pink) hydrological connectivity.

| Type | Description | Viability of passage for eels, and justification | n |
| --- | --- | --- | --- |
| 0 | None | Very low: no known route | 24 |
| 1 | Pumped water transfers | Low: flow direction is not attractive to upstream moving eels and pump mechanism must be safely navigated | 3 |
| 2 | Non-pumped water transfers | Moderate: flow direction is not attractive to upstream moving eels | 96 |
| 3 | Low banks known to flood | Moderate-high: flooding is a natural fish dispersal mechanism (Bolland et al., 2015), but may be sporadic | 9 |
| 4 | Gravity (non-pumped) drainage | High: flow direction is attractive to upstream moving eels (Piper et al., 2012) | 17 |

Habitat management scores were derived from the frequency (i.e. never, less than every 5 years, every 1-5 years, yearly and more than yearly) of bank trimming, in-channel weed cutting, desilting and dredging. Both weighted and unweighted additive (sum) and multiplicative (product) scores were calculated. Unweighted scores assumed all four habitat management methods were equally influential on the prevailing fish community, and that increasing frequency of each method was more detrimental (Table S2). The weighted scores assigned increased weighting to practices deemed more detrimental to habitat: dredging was deemed most detrimental, followed by desilting, followed by in-channel weed removal, while bank trimming was considered the least detrimental (Table S3). These weighted scores were also combined to produce additive and multiplicative management indices.

Fish species richness was calculated as the total number of fish taxa detected per catchment across all samples. When comparing sites with and without eels present, European eels were removed from species richness. Observed species richness was used for analyses that explicitly accounted for sampling effort, such as modelling inter-round effects on species richness as all sites had the same number of samples taken. To account for variation in sampling effort due to the removal survey design, fish species richness was standardised to one sample per site using interpolation with iNEXT and referred to hereafter as single-sample species richness. At sites with only a single sample, observed richness was used directly. Rheophilic and marine fish species were defined using flow guilds from Aarts & Nienhuis (2003) with the exception of spined loach (*Cobitis taenia*) which were reclassified from rheophilic to eurytopic due to their now-known abundance in drain habitats (Nunn et al., 2014).

Sites were administered by 46 local Internal Drainage Boards (IDBs), which are responsible for the day-to-day operation and maintenance of small groups of pumping stations. These local IDBs fell under six wider drainage authorities. Differences in single-sample species richness were tested among drainage authorities (n = 6) to provide a meaningful number of groups for analysis. Conversely, local IDB was used to test for habitat management clustering and included as a random intercept when modelling single-sample species richness to account for spatial clustering and the homogeneity of local habitat management practices.

### Statistical analyses

The level of confidence in absence at eel-negative sites according to the number of samples taken was quantified using the confidence in absence for decision making (CIADM) model developed by Griffiths et al. (2025b). We used the PCR replication (*P_r_*) dataset from Griffiths et al. (2025b) to inform our confidence estimations, given it was for European eel upstream of pumping stations, as is under investigation here.

Where data did not meet the assumptions of normality (tested with Shapiro-Wilk test, inspected visually with histograms), non-parametric tests were used. Differences in single-sample species richness between eel-positive and eel-negative sites were tested using Mann–Whitney U tests. Spearman’s rank correlation coefficients were used to assess associations between continuous variables, i.e. distance from tidal limit and catchment area. Fisher’s exact tests were used to assess associations between categorical variables, i.e. presence of individual species (e.g. spined loach), groups of species (i.e. rheophilic, marine), hydrological connectivity, number of barriers and the frequency of management practices. Kruskal–Wallis tests were used to assess differences in continuous variables among groups with more than two categories: single-sample species richness, distance from the tidal limit and catchment area among hydrological connectivity groups, and single-sample species richness across habitat management frequencies. Where Kruskal–Wallis tests indicated significant differences, post-hoc pairwise comparisons were conducted using Dunn’s tests with Benjamini–Hochberg correction for multiple comparisons. Cramér’s V statistics were used to assess clustering of habitat management frequencies at sites by owned by different local IDBs. All tests were conducted using a minimum significance threshold of 0.05.

European eel presence against abiotic (i.e. distance from tidal limit, catchment area, habitat management and hydrological connectivity) and biotic (i.e. fish species richness) variables was modelled using generalised linear models (GLMs) with a binomial error distribution and logit link. The habitat management scores were modelled as continuous variables. The type of hydrological connectivity (Table 1) was tested individually (i.e. binary form), in combination (i.e. all five types) and in groups (i.e. natural, manual and none) as categoric variables. The source of water for sites with manual connectivity (neighbouring catchment or downstream waterbody) was also tested. Model selection was based on lowest Akaike’s Information Criterion (AIC) and significance of effects (p<0.05) and performance was assessed using McFadden’s pseudo-R². Model diagnostics included visual inspection of residual plots, assessment of overdispersion using Pearson residuals, and examination of leverage and influence using Cook’s distance. Separation was tested and variance inflation factors were used to confirm absence of problematic multicollinearity.

The effects of abiotic variables (i.e. distance from tidal limit, catchment area, habitat management and hydrological connectivity) on single-sample species richness were modelled using a Gamma generalized linear mixed model (GLMM) with a log link. The source of water for sites with manual connectivity (neighbouring catchment or downstream waterbody) was also tested as a categoric variable. Manual sites with water transferred from the downstream waterbody were also combined with Type 4 sites, i.e. those with a gravity sluice, and were treated as a binary variable against all other sites. Habitat management scores were handled the same as the European eel presence GLM. Local IDB was included as a random intercept to account for spatial clustering of sites. Model assumptions were assessed using simulated residuals; uniformity, dispersion and outliers were tested and multicollinearity among fixed effects was assessed using variance inflation factors.

The influence of temporal variations in hydrological connectivity (same groups as previous model) on all-sample species richness were also modelled (GLMM) using Poisson mixed effects with site included as a random effect to capture site-level changes. The models used the first two samples from rounds 2 (spring), 3 (summer)and 4 (early autumn) at sites that were sampled in all three of those rounds (n = 90) to enable fair comparison of fish species detection. Round 1 was excluded because 17.1% of sites (n = 26) were inhibited (i.e. fish detection in these samples was unreliable) and 93.4% of sites (n = 142) only had a single sample taken. For details of handling of inhibited samples, see Monaghan et al. (2026). Model adequacy was evaluated using standard mixed-model diagnostic procedures, including checks for singularity, overdispersion, simulated residual diagnostics, residual plots, collinearity, influential sites, and random-effects distributions. Fish assemblage composition was analysed using non-metric multidimensional scaling (see supplementary material).

All statistical analyses were carried out in R version 4.5.1 (R Core Team, 2025) with data processing performed using the tidyverse (Wickham et al., 2019) package and analyses utilising iNEXT (Hsieh et al., 2020), glmmTMB (Brooks et al., 2017), vegan (Oksanen et al., 2025), detectseparation (Kosmidis et al., 2026), pscl (Jackman et al., 2024), ggeffects (Lüdecke et al., 2025), performance (Lüdecke et al., 2026) and DHARMa (Hartig et al., 2026) packages. Mapping was carried out using a custom workflow in R utilising sf, maps and mapdata packages.

## Results

### Landscape-scale eel distribution

European eels were detected in 43 of the 152 (28.3%) pumped catchments sampled; one of one (100%) that discharges into the sea, nine of 15 (60.0%) that discharge into an estuary and 33 of 136 (24.3%) that discharge into fresh water (Figure 2). Five, 53, 25 and seven eel-negative sites were sampled five, seven, eight and 10 times, respectively, which corresponded to >92%, >96%, >97% and >98% confidence in eel absence (Figure 2). Sites with a single sample in R1 that were screened out for poor quality habitat still had a confidence in absence of >74%.

**Figure 2.**
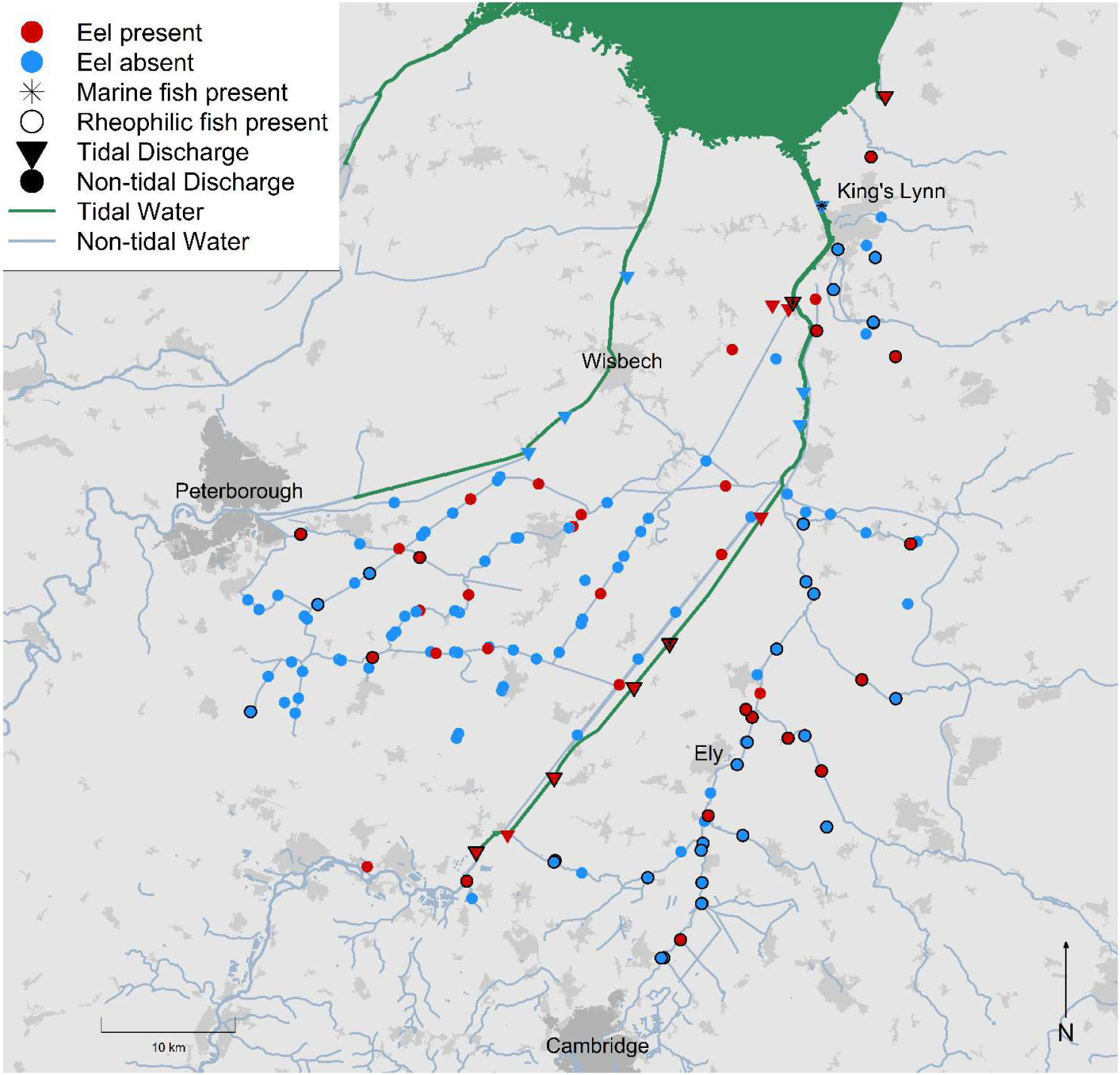
Pumping station catchments in The Fens that were identified using eDNA metabarcoding as being European eel positive (red) and negative (blue). Urban areas (dark grey) and major tidal (green) and non-tidal (blue) water bodies are marked. Presence of rheophilic fish species (black outlines) and marine fish (*) are marked. Built-up area and settlement data: Office for National Statistics, licensed under the Open Government Licence v3.0. Contains OS data © Crown copyright and database rights 2026. Waterway data © OpenStreetMap contributors.

### Abiotic predictors of eel occurrence

In the abiotic eel model, eel occurrence declined significantly with increasing distance from the tidal limit (β = −0.38 ± 0.15 SE, p = 0.011) but increased where natural connectivity (Type 3, flooding and Type 4, gravity sluices) was present (β = 1.30 ± 0.48 SE, p = 0.007). Eel presence was also positively associated with catchment area (β = 0.46 ± 0.17 SE, p = 0.007). Importantly, there was no difference in catchment area between sites with different types of hydrological connectivity (Kruskal-Wallis test, X^2^ = 3.9003, df = 2, p = 0.1423). This abiotic model explained approximately 15% of the deviance (McFadden’s pseudo-R² ≈ 0.149) (Figure 3).

**Figure 3.**
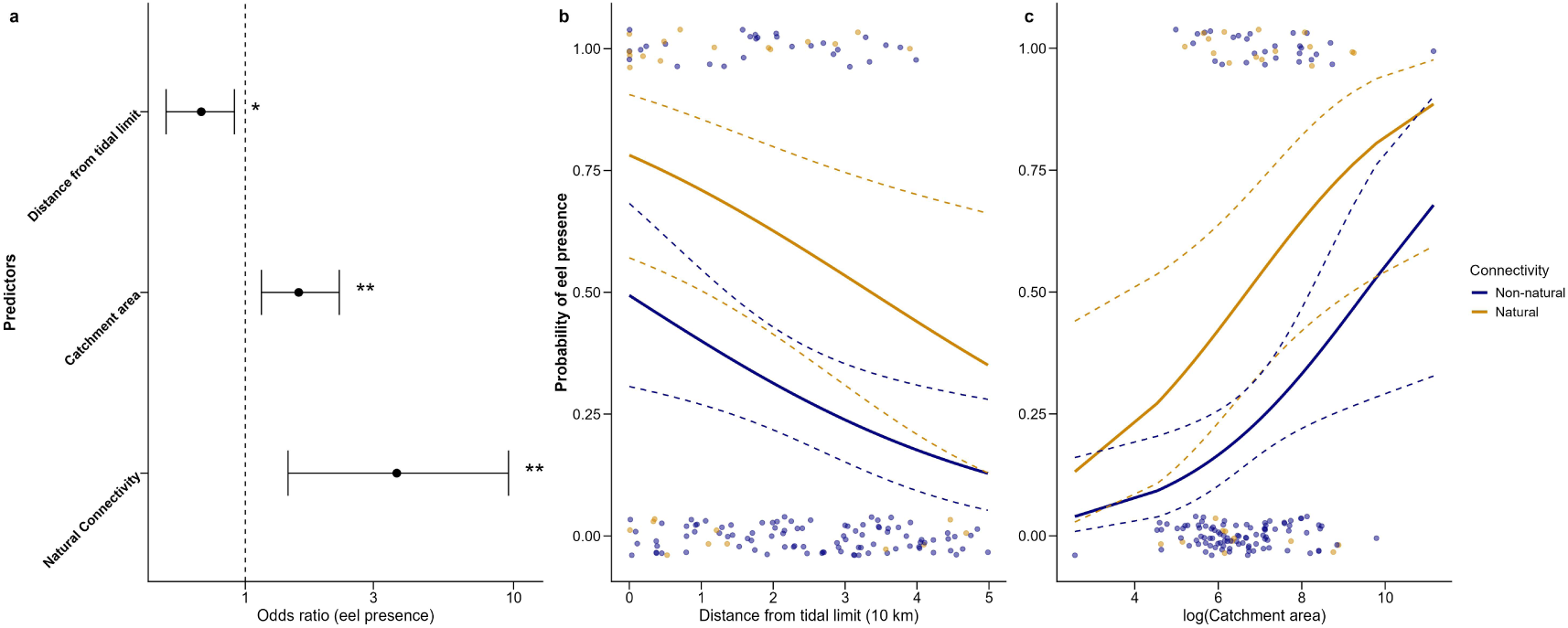
Abiotic eel detection model odds ratios (a) and probabilities of eel detection in relation to tidal distance (b) and catchment area (c) for natural (Types 3 and 4) and non-natural (Types 0, 1 and 2) hydrological connectivity groups (Table 1). Dashed lines denote 95% confidence intervals. * and ** denote 95% and 99% significant differences, respectively. See Figure S1 for the distance from the tidal limit and catchment area for sites within each to hydrological connectivity group.

European eels were not detected upstream of 88 of the 125 (70.4%) pumping stations with known hydrological connectivity to downstream waterbody or neighbouring catchment (Types 1-4). Specifically, eel were detected in 58.8% (10 of 17) of gravity sluice sites (Type 4) and 55.6% (five of nine) of sites with over-bank flooding (Type 3), i.e. those with natural hydrological connectivity, and significantly positively influenced eel presence (Fisher’s exact test, p = 0.0007; odds ratio = 4.57, 95% CI: 1.74–12.40, Figure 3a). Conversely, manual water transfers into the catchment (22.2% eel-positive; 22 of 99) showed a weaker, negative association (p = 0.0208; odds ratio = 0.40, 95% CI: 0.18–0.88). Distance from the tidal limit differed significantly across connectivity types (Kruskal–Wallis test: X² = 30.15, df = 2, p < 0.001) (Figure S1). Post hoc Dunn tests with Benjamini–Hochberg correction showed there was no significant difference detected between manual and natural connectivity types (adjusted p = 0.159). By contrast, disconnected (“none”) sites were located significantly closer to the tidal limit than both manually and naturally connected sites (adjusted p < 0.001 for both comparisons). Despite this, European eels were only detected at 25.0% (six of 24) of sites with no reported present-day hydrological connectivity (Type 0). Eel occurrence was also negatively impacted by the number of downstream barriers (Fisher’s exact test, p = 0.003), with eels detected at 66.7% (n = 15), 28.2% (n = 103), 14.3% (n = 28) and 0% (n = 3) of sites upstream of 0, 1, 2 and 3 barriers, respectively.

Neither individual (Fisher’s exact test, all p >0.05) nor combined (weighted and unweighted, additive and multiplicative; Mann-Whitney U test, all p > 0.6) habitat management practices had a significant influence on eel occurrence. All physical habitat maintenance activities significantly clustered across drainage authorities (X^2^; p < 0.001 in all cases); the strongest association was for dredging (Cramér’s V = 0.98), followed by desilting (Cramér’s V = 0.93), in-channel weed cutting (Cramér’s V = 0.92) and bank trimming (Cramér’s V = 0.83). Despite this, the likelihood of eel occurrence did not differ significantly across wider drainage authorities (n = 6) (Fisher’s exact test, p = 0.084). Incorporating each habitat management score into a model of eel presence in place of either catchment area or single-sample species richness produced non-significant effects (p all >0.5, Figure 4).

**Figure 4.**
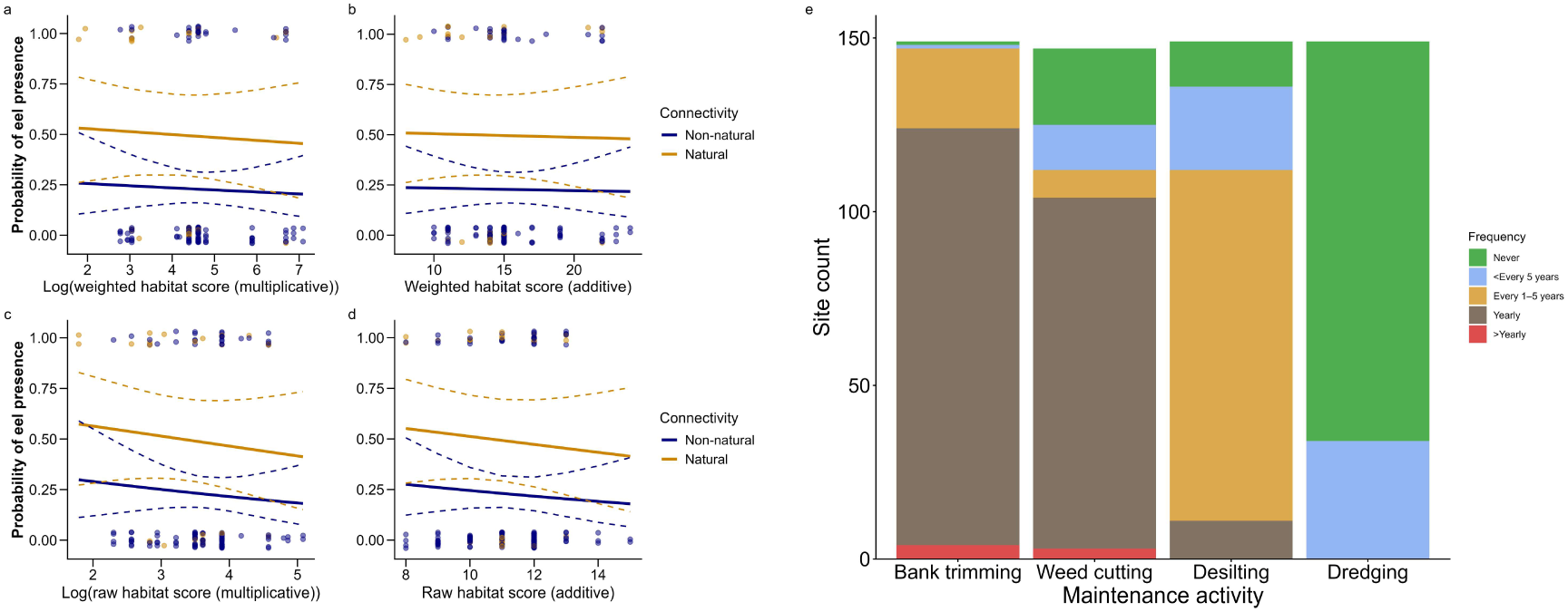
Probability of eel occurrence upstream of pumping stations with natural (Types 3 and 4) and non-natural (Types 0, 1 and 2) hydrological connectivity (Table 1) in relation to weighted and un-weighted habitat management scores (a-d), and the frequency of each of the management activities in pumped catchments in The Fens (e).

### Incorporating a biotic predictor of eel occurrence

Eel occurrence probability increased significantly with increasing single-sample species richness (β = 0.41 ± 0.11 SE, p < 0.001) modelled in place of catchment area, while the presence of natural migration routes remained positively associated with eel occurrence (β = 1.17 ± 0.52 SE, p = 0.023). The probability of eel occurrence also retained a significant negative relationship with increasing distance from the tidal limit (β = −0.62 ± 0.18 SE, p < 0.001). This biotic model explained approximately 21% of the deviance (McFadden’s pseudo-R² = 0.212) (Figure 5).

**Figure 5.**
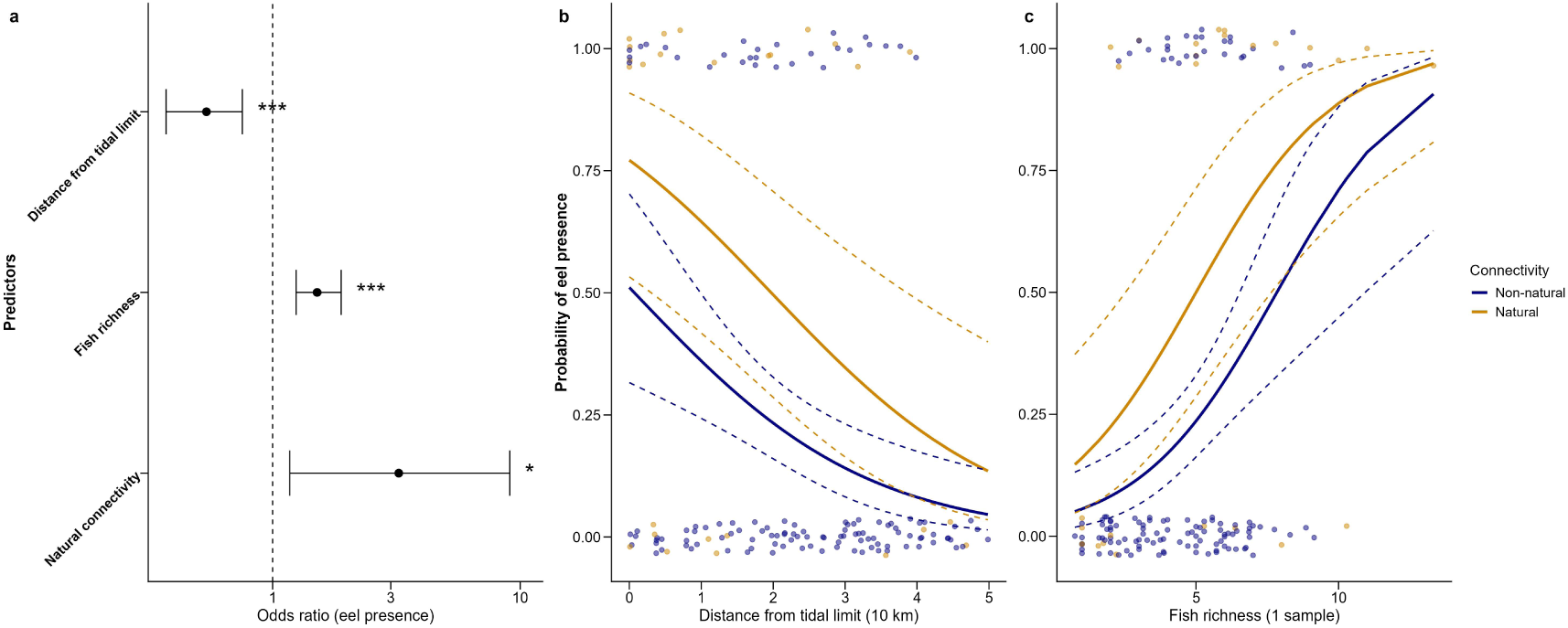
Biotic eel detection model odds ratios (a) and probabilities of eel detection in relation to tidal distance (b) and single-sample species richness (c), split by natural and (Types 3 and 4) and non-natural (Types 0, 1 and 2) hydrological connectivity (Table 1). Dashed lines denote 95% confidence intervals.

The occurrence of marine or rheophilic fishes was also explored as a proxy for upstream connectivity for eels; marine fish species were detected upstream of three of 16 pumping stations at the tidal limit, one of which was negative for eel (Figure 2; Figure S2). Rheophilic fish species were detected upstream of 43 of 136 pumping stations that discharge into fresh water (Figure 2; Figure S2) and median rheophilic fish species richness was higher at eel-positive (1 (IQR = 0-1)) than eel-negative sites (0 (IQR = 0-0)) (Mann-Whitney U test, W = 1755, p = 0.004). Notwithstanding, rheophilic fish species were detected at 27 of 109 (24.8%) sites that were negative for eel; 20 of 81 (24.7%) with manual water transfers (Types 1 and 2), two of seven (28.6%) with a gravity sluice (Type 4), five of 18 (27.8%) with no known hydrological connectivity (Type 0) and one of four (25.0%) with flooding (Type 3) and zero of 3 (0.0%) of sites with unknown connectivity. Rheophilic-positive sites were also significantly further from the tidal limit than eel-positive sites (Mann-Whitney U test, W = 789, p = 0.01209).

### Predictors of fish species richness

Catchment area was positively associated with single-sample species richness (effect ratio = 1.20, 95% CI: 1.12–1.28, p < 0.001). Sites with manual transfers from the downstream water body (Types 1 and 2) or gravity sluices (Type 4) supported significantly higher richness than other sites (effect ratio = 1.37, 95% CI: 1.13–1.67, p = 0.002) (Figure 6). A random intercept was included for IDB to account for spatial and habitat management clustering. The model explained a moderate proportion of variation in single-sample species richness (marginal R² = 0.22, conditional R² = 0.54).

**Figure 6.**
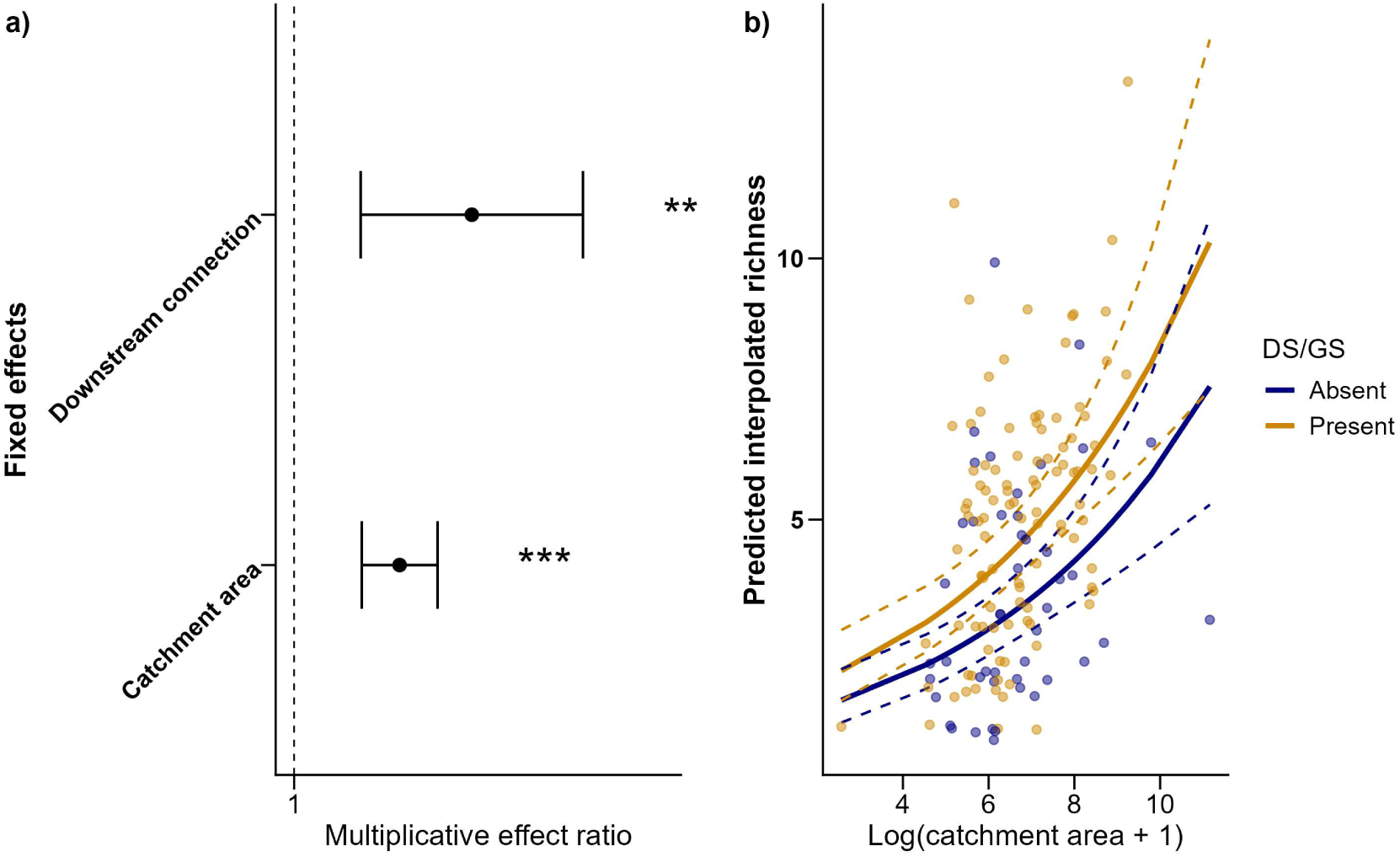
Effect ratios for all predictors in the single-sample species richness model (a) and in relation to catchment area, split by the hydrological connectivity (Types 1 and 2 from the downstream waterbody or Type 4(Table 1); labelled “DS/GS”) (b).

The median single-sample species richness was significantly higher at eel-positive (5.20, IQR = 4.07–6.83) than eel-negative sites (3.86, IQR = 2.00–5.86) (Mann–Whitney U test, W = 1452.5, p < 0.001) (Figure 7). The median single-sample species richness was significantly different among hydrological connectivity groups (Kruskal–Wallis X² = 13.33, p = 0.001). Sites with no hydrological connectivity (Type 0) had significantly lower richness than both manually (post hoc Dunn’s test with Benjamini-Hochberg correction; p = 0.0004) and naturally (p = 0.0105) connected sites. There was no significant difference between manual and natural connectivity (p = 0.2711) (Figure 7).

**Figure 7.**
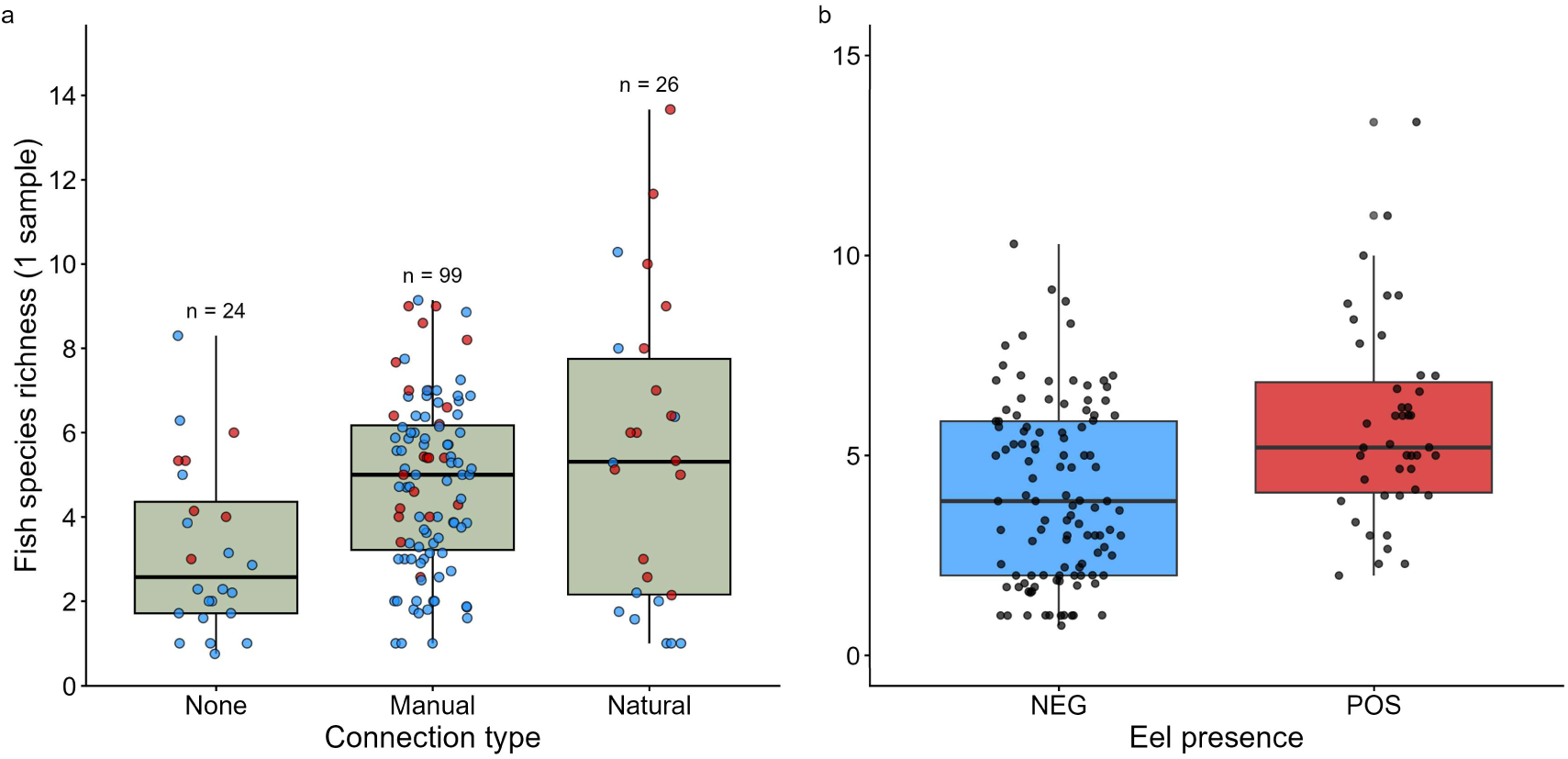
Single-sample species richness according to hydrological connectivity type (Table 1) (a) and eel presence (b). Red and blue points (a) denote sites with and without eels detected upstream, respectively.

At sites with manual water transfers from the downstream water body, observed fish species richness was significantly higher during R3 (summer) (incidence rate ratio (IRR) = 1.22, p = 0.010) and R4 (early autumn) (IRR = 1.30, p < 0.001) compared to R2 (spring). At sites without manual transfers from the downstream water body, there was no significant variation in richness across visits (R3: IRR = 1.00, p = 1.000; R4: IRR = 1.05, p = 0.689) (Figure 8). Overall, there was limited separation of fish assemblages between eel-positive and eel-negative sites across sampling rounds (see supplementary material; Figure S3).

**Figure 8.**
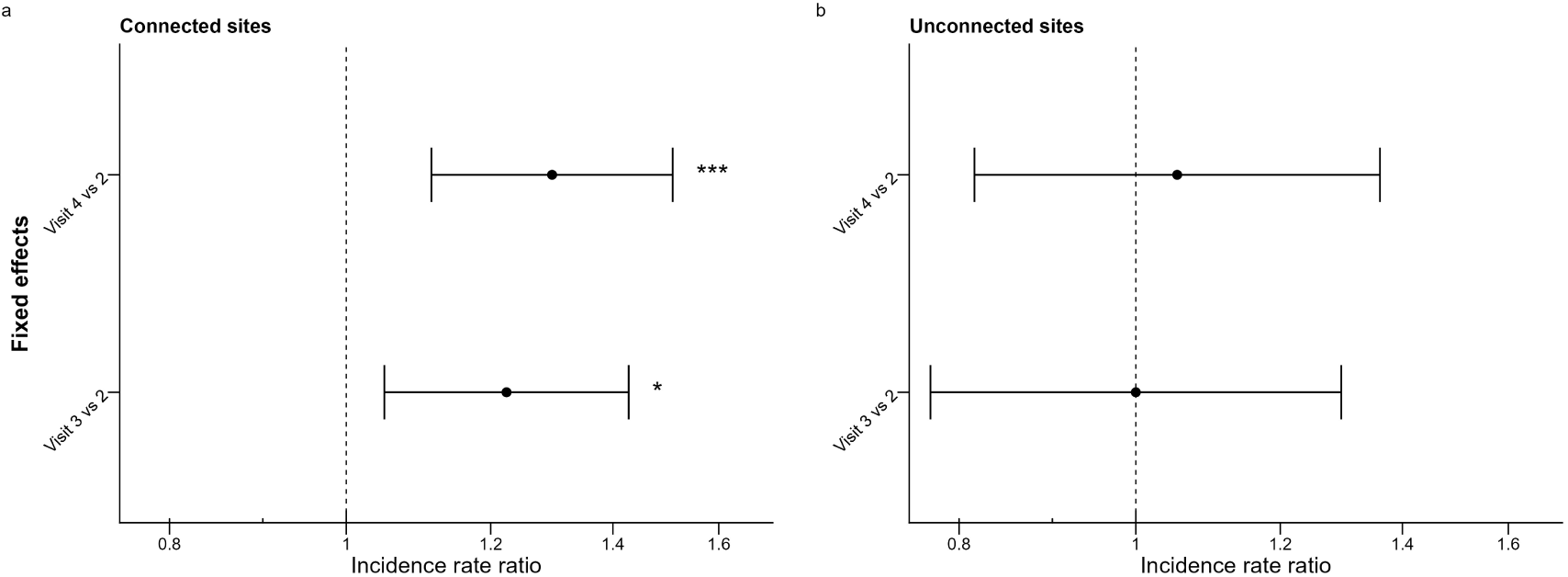
Estimated incidence rate ratios (IRR ± 95% CI) for all-sample species richness during R3 (summer) and R4 (early autumn) relative to R2 (spring) at sites with (a) and without (b) manual water transfers from downstream. * and *** denote 95% and 99.9% significant differences, respectively.

There was no significant difference in median single-sample species richness where single and double bank flail mowing took place (Mann-Whitney U test, W = 2774.5, p = 0.8901). None of the physical habitat scores were correlated with single-sample species richness (Spearman’s rank, rho = 0.0-0.06, p >0.48). Single-sample species richness did not differ across individual maintenance variable frequencies (Kruskal-Wallis test, all p >0.3) aside from the frequency of in-channel weed cutting (Kruskal-Wallis test, X^2^ = 16.672, df = 4, p < 0.005). A post hoc Dunn test with Benjamini–Hochberg correction showed that a frequency of “every 1-5 years” (n = 8) had significantly lower richness than all other frequencies (adjusted p-values = 0.003-0.017).

## Discussion

There is an urgent need to establish the present-day distribution of rare and elusive species, especially in anthropogenically dominated landscapes with implications for species conservation as well as infrastructure and catchment management. Here, we establish and assess landscape-level distribution of critically endangered European eel upstream of pumping stations, given legislative drivers, in a region subject to considerable land-use change. Once an extensive wetland, The Fens have been intensively drained and are now characterised by intensive agriculture practices and heavily managed drainage channels. Wider fish community data was also incorporated as biotic indicators to aid understanding of ecological processes and impacts of fragmentation, variable hydrological connectivity and habitat quality. Specifically, eels were detected upstream of 28.3% of the pumping stations studied, with higher rates of detection in significantly larger catchments, stations closer to tidal waters and the presence of natural hydrological connectivity (gravity sluice or overbank flooding) being abiotic predictors of eel distribution.

The present-day distribution of European eel upstream of pumping stations in The Fens was established using eDNA, a highly sensitive method for detecting eels in pumped catchments (Griffiths et al., 2020), with a high confidence in absence (>95%; Griffiths et al., 2025) using a removal survey design (Mackenzie & Royle, 2005). Historically, eels were highly abundant across the pre-drainage wetlands in The Fens landscape (Porter, 1958) but the present-day occupancy of pumped catchments was low (28.3%). This is consistent with previous investigations into eel presence in pumped catchments, with Griffiths et al. (2025a) reporting occupancies below 10% in pumped catchments in the River Ancholme, England. Low European eel site occupancy has also been found in other anthropogenically impacted landscapes, including highly fragmented rivers in Cyprus (Griffiths et al., 2023) and hydroelectric power stations in Norway (Halvorsen et al., 2020). Similar has also been reported for American eel (*Anguilla* rostrata) at navigation locks in North America (George et al., 2023) and significant reductions in freshwater distributions of tropical Anguillid eels in South Africa and Mozambique (Hanzen et al., 2022). Overall, the significant decline in European eel has also culminated in reductions in distribution, which in this case was exacerbated by land use change and anthropogenic reductions in connectivity.

Pumping stations and associated infrastructure (e.g. flood banks) fragment longitudinal and lateral hydrological connectivity. Here, significant abiotic predictors of eel detection were being closer to tidal waters, having larger catchments and the presence of either a non-pumped gravity drainage route (Type 4) or hydrological connectivity during over-bank flooding (Type 3). Eel presence in sites closer to the tidal limit is logical given their catadromous life cycle (Schmidt, 1923) and, coupled with probability-driven dispersion due to greater numbers of juveniles present in tidal waters (Itakura & Wakiya, 2020), their ingress into pumped catchments near tidal waters would be more likely. Once in freshwater, European eel tend to be larger and in lower densities further inland (Naismith & Knights, 1993; Laffaille et al., 2003); density-driven dispersal is documented in eels (Bevacqua et al., 2011). They also have high habitat fidelity and small home ranges (Baras et al., 1998; Laffaille et al., 2003; Ovidio et al., 2013; Verhelst et al., 2018). All these factors could culminate in eels being less likely to attempt passage into pumped catchments further from the sea. Increases in numbers of barriers further inland also contributed to lower site occupancy, as has been documented elsewhere (Halvorsen et al., 2020; Griffiths et al., 2023; Macarthur et al., 2025). Overall, the significant negative effect of tidal distance on eel distribution at such a small resolution (all sites within 100 km of the tidal limit) is likely a result of eels’ extreme decline and landscape scale fragmentation.

This investigation uniquely established, at a landscape scale, which types of hydrological connectivity significantly affect eel distribution. Twenty-four of 152 pumping stations across The Fens were completely isolated hydrologically (when pumps were not operating; Type 0), but European eels were detected in six of these pumped catchments. This may reflect the well-established ability of European eel to climb wetted surfaces (Podgorniak et al., 2016; Piper et al., 2023) and thus may have crawled up pumping station pipework or through leaks in the infrastructure. Alternatively, it may reflect a remanent population from historic connectivity, as has been reported upstream of dams (Podda et al., 2022). All other pumping stations catchments in The Fens were categorised according to the type of hydrological connectivity with the downstream/receiving waterbody, i.e. downstream gravity discharge (Type 4), over-bank flooding (Type 3) and upstream manual transfers (both pumped (Type 1) and non-pumped (Type 2)). The first two, in combination, represent natural hydrological connectivity and was a significant positive predictor of eel presence. We speculate this corresponds to the migratory propensity of glass eel and elver life-stages to migrate upstream into rivers (Laffaille et al., 2007; Kroes et al., 2020; Monteiro et al., 2023), while acknowledging subtle variation in hydrodynamic conditions at human-made barriers can influence upstream passage (Piper et al., 2012). Upstream movements of yellow-phase shortfin eel (*Anguilla australis*) has also been demonstrated at a flood relief pump station with gravity bypass in New Zealand (Mahlum et al., 2025). Lasne et al. (2008) reported European eel colonisation of laterally disconnected floodplain waterbodies in the River Loire, France, during floods. By contrast, sites that transfer water in an upstream direction into the pumped catchment had reduced eel presence, potentially because the direction of water flow was misaligned with the migratory tendencies of both juvenile and yellow European eel.

Upstream manual transfers of water into pumped catchments in The Fens is typically performed to maintain water levels during agricultural irrigation. Stations with upstream manual transfers were a comparable distance from the tidal limit as stations with gravity sluices, which has implications for the timing of hydrological connection for eel movements into pumped catchments. Manually operated penstocks were the dominant transfer mechanism (86.9%), which will predominantly occur during the working day, and irrigation tends to occur during mid-to-late summer. Consequently, such upstream water transfers do not align with diel peaks in eels activity, being almost exclusively nocturnal or crepuscular (Barry et al., 2016; Verhelst et al., 2018), nor seasonally with their typical habitat colonisation period in winter/spring (Moura et al., 2022; Boardman et al., 2023). Furthermore, rheophilic fish species were detected in 24.7% of the eel-negative sites with manual connectivity, thus demonstrating they are viable fish passage routes. Elsewhere water transfers for irrigation purposes have been shown to negatively impact fish populations via displacement and cause significant mortality (King & O’Connor, 2007; Boys et al., 2021).

Regardless of immigration route, the habitat in a pumped catchment must be of sufficient quality for eels to survive until they emigrate, either as yellow eels moving into habitat further downstream or as silver eels during their seaward migration to the Sargasso Sea, potentially up to 30 years later (Durif et al., 2020). Pumped catchments are characterised by extensive and intensive physical habitat maintenance measures to ensure water conveyance during flood-relief pumping (Ward-Campbell et al., 2017; Guo et al., 2024), but they were not a predictor of European eel distribution, neither in isolation or in combination. This could be attributed to the generalist nature and wide tolerance ranges for physical habitat and water quality conditions (Townsend & Peirson, 1988; Daverat et al., 2006; Pujolar et al., 2012; Pike et al., 2020; Enbody et al., 2021; Näslund et al., 2022), the lack of variation in the type and frequency of river maintenance between sites and the confounding influence of connectivity. Alternatively, drain habitat management practices may influence European eel population structure, condition and abundance, rather than presence/absence, and thus would not be detectable with eDNA. Indeed, studies have demonstrated individual fish mortality, as well as changes in fish density and community composition, associated with benthic substrate and macrophyte disturbance and removal (Freedman et al., 2013; Thiemer et al., 2021). However, resilience of fish communities to drain management has also been documented (Ward-Campbell et al., 2017) and it could also be argued that removal of weed and anoxic silt from excessively eutrophic drains could be beneficial to fish assemblages by helping to increase oxygen levels (Zarull et al., 2002; Perna & Burrows, 2005; Mitsuo et al., 2014).

In the abiotic model, eel positive sites had significantly larger catchment area, and thus catchment area may be a proxy for habitat quality. Catchment area in the abiotic model was replaced by fish species richness in the biotic model and had superior performance (21.2% variance explained rather than 14.9%). This suggests the prevailing fish community was a long-term biotic indicator of habitat quality (Karr, 1981; Blabolil et al., 2017). Furthermore, significantly larger catchments were predictors of fish species richness. Indeed, larger catchments may have more suitable habitat, e.g. wider and deeper channels, or increased habitat heterogeneity, which may be required for feeding, as well as flow and predator refuge (Beecher et al., 1988; Kärnä et al., 2019; Beevers et al., 2021). Fish communities in larger catchments may also be more resilient than those of smaller catchments by increasing the potential to escape harmful events such as pollution, eutrophication, extreme heat and dissolved oxygen crashes (Caissie, 2006; Wollheim et al., 2022). Small and heavily managed pumped catchments with uniform in-channel and riparian habitat are devoid of flow and predator refuge (Norman et al., 2024) and may be vulnerable to poor water quality and thermal events during heatwaves (Rideout et al., 2022) as well as winter freezing. Hydrological connectivity to the downstream/receiving waterbody likely confounds this by enabling fish movement into the pumped catchment. Indeed, gravity sluices and manual transfers from the downstream water body were a significant positive predictor of fish species richness. Gravity sluices can enable open connection (e.g. pointing doors) and thus the fish community is effectively an extension of the downstream river channel rather than an isolated sub-catchment. Catchments with regular connectivity could also be quickly recolonised by fish despite having habitat that is either sub-optimal or not resilient to environmental stressors (De Miguel et al., 2016; Kristensen et al., 2020; Sun et al., 2022; Kiffney et al., 2023).

While good hydrological connectivity may increase recolonisation and recovery following disturbance, it may, however, perpetually transfer fish into vulnerable habitats. Significantly disproportionate increases in fish richness were found at sites with manual transfers from the downstream water body during summer sampling (R3 and R4), likely due to the occurrence of upstream transfers for irrigation at this time. While it could be argued that water transfers into a pumped catchment may transfer fish eDNA rather than actual fish, unscreened inlets that transfer large quantities of water must be treated as a potential route for fish movement, particularly for small and/or juvenile fishes. Similar findings were not reported in catchments that do not transfer from the downstream water body, and thus are unlikely to be attributed to seasonal variations in eDNA detection (Griffiths et al., 2025a). Fish that enter during this time must persist once transfers cease and the catchment becomes isolated. Given the impoverished fish community at other times of year, we speculate this is not the case, and sampling outside of peak irrigation season yields a truer reflection of the prevailing habitat and water quality. Overall, pumped catchments with manual water transfers are potentially ecological traps given transfers from the downstream water body may replenish fish populations that are unable to persist during periods of disconnection. Artificial water bodies with modified hydrological connectivity elsewhere have also been demonstrated as ecological traps for fish (Pelicice & Agostinho, 2008; Hale et al., 2018).

The most site-abundant fish species detected are consistent with those typically associated with lentic conditions typically found in fenland drainage habitat (Townsend & Peirson, 1988; Aarts & Nienhuis, 2003). The presence of marine and rheophilic fish species which require flow for reproduction (Aarts & Nienhuis, 2003) were also used as biotic indicators of upstream passability into the pumped catchments. Elsewhere, rheophilic species, such as stone loach, dace, gudgeon and chub, are known to inhabit still or low flow waters like canals, ponds and lakes, provided there is connectivity to lotic waters (Schiemer & Waidbacher, 1992; Arlinghaus & Wolter, 2003; Hänfling et al., 2016). From an eel perspective, they help further understanding of where the hydrological connection is not conducive to upstream eel passage. Importantly, much like European eel, marine and rheophilic fish must be able to safely return to the downstream waterbody to complete their lifecycle (Aarts & Nienhuis, 2003).

Overall, the use of eDNA sampling allowed us to establish eel and resident fish distributions with higher sensitivity and reduced costs, field effort and invasiveness of sampling compared to those of traditional fish sampling methods (Griffiths et al., 2020). Indeed, over 855 eDNA samples were collected from 152 sites at landscape-scale during four sampling campaigns over two and a half years, and thus represents one of the most extensive and comprehensive eDNA investigations to-date globally. However, further eDNA studies in pumped catchments could employ sampling both upstream and downstream of pumping stations to truly assess their impact as a barrier to eel and resident fish populations. The presence/absence data provided by eDNA does not facilitate comparison of eel density or individual fish (e.g. lengths, weights, body condition, life-stage) across catchments. Therefore, potential further research could look to use capture-based surveys in the eel-positive catchments to compare size, structure and condition of populations.

## Conclusion

This study established the distribution of critically endangered European eel in a historic wetland landscape now characterised by intensive arable farming and heavily managed drainage channels. The biotic model had improved and strong explanatory performance for a binary ecological response, with species richness replacing catchment area in the abiotic model, which we speculate is attributed to fish being improved indicators of connectivity, physical habitat quality and water quality. Catchments with eels detected also had significantly richer fish communities and rheophilic fish species than those without. Manual upstream water transfers for irrigation were a genuine route for fish to enter a catchment but not European eel, likely attributed to nuances associated with species ecology coupled with flow direction and timing of inlet operation. Notwithstanding, if eel populations recover significantly in the future, density driven competition for habitat could increase likelihood of eels entering pumped catchments through irrigation inlets. In the meantime, the findings have implications for species conservation and infrastructure management, including mechanisms to protect eels while both inhabiting and emigrating from pumped catchments (Baker et al., 2019; Evans et al., 2024b, 2025; Carter et al., 2025) to comply with legislation (EC Regulation No. 1100/2007). At sites without eels, efforts could include improving immigration, provided the reason for absence is fully understood and can be mitigated. Ultimately, given the socioeconomic reliance on pumping stations in The Fens, the landscape cannot be restored to its pre-drainage state but efforts to conserve critically endangered European eel and improve catchments for entire fish communities without compromising drainage efficiency should be pursued.

## Declarations

## Acknowledgments

We thank the Environment Agency for funding the study and drainage authority staff for granting site access and assisting with data collection, particularly C. Hipkin, R. Taylor, C. Laburn, L. Butler, A. Sweeney and E. Johnson. We thank C. Cowgill, C. Collins, R. Donnelly and O. Evans for assistance with sample collection and laboratory work. We thank D. Pollard for allowing access to Denver Sluice Complex. We thank K. Jerrom for input when devising the project.

## Author Contributions

Sample collection was carried out by J.D.B. and A.I.T.M. Laboratory work was carried out by A.I.T.M, N.P.G, J.A.M and G.S.S. Data analysis was carried out by A.I.T.M, N.P.G, G.S.S, and M.C. Bioinformatics analyses were completed by G.S.S. The study was conceived and designed by J.D.B, R.M.W, N.P.G, A.D.N, L.L.H, and B.H. Funding was acquired by J.D.B, R.M.W, and B.H. The initial manuscript draft was prepared by A.I.T.M. All authors contributed to manuscript review and approved the final version.

## Data availability

Data and scripts have been archived and made available on Zenodo. Raw eDNA outputs: https://doi.org/10.5281/zenodo.19099273, data analysis: https://doi.org/10.5281/zenodo.22087768.

