## Supplementary material for "Pumping stations negatively affect the distribution of critically endangered European eel (*Anguilla anguilla*); a landscape-scale study using environmental DNA metabarcoding"

### Methods

#### Hydrological connectivity

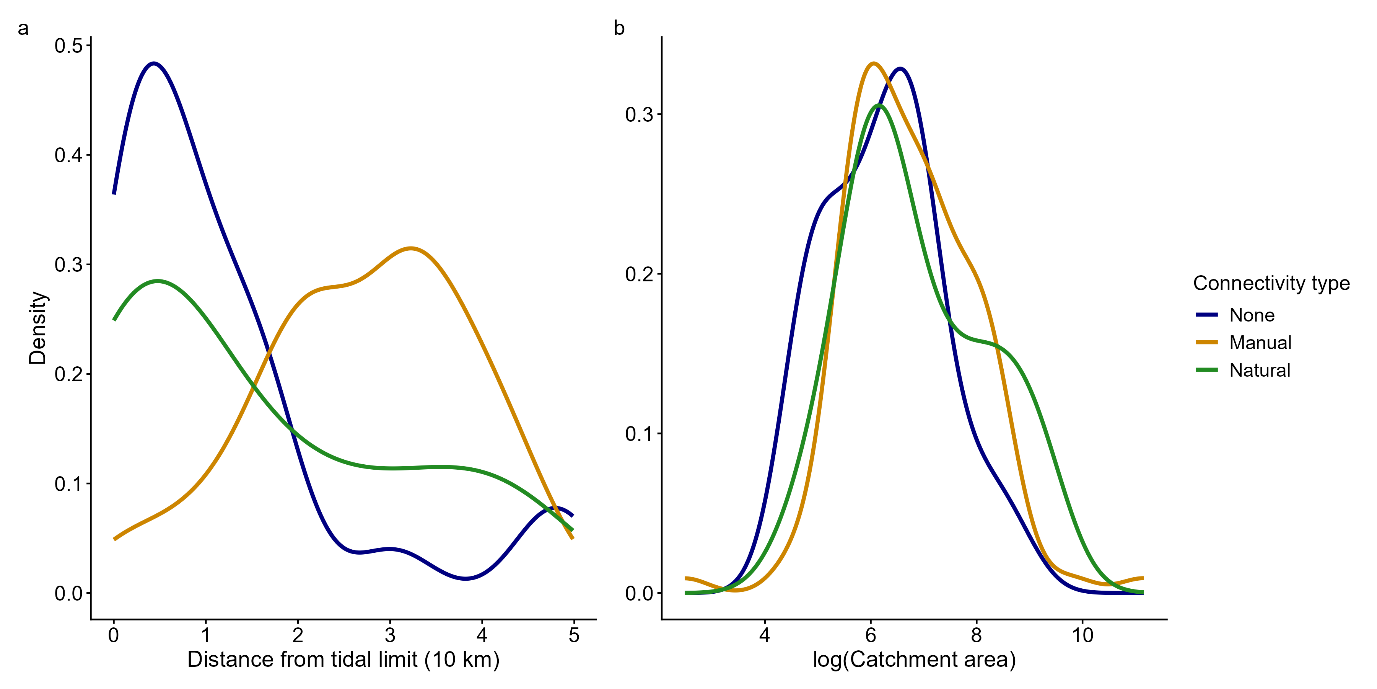

Figure S1. Distributions of pumping station with none, manual and natural hydrological connectivity (Table 1) in relation to (a) distance from tidal limit and (b) catchment area.

Table S1. Frequencies and methods of sites that manually transferred water

| **Frequency** | **Pumped** | **Sluice/penstock** | **Other** |
| --- | --- | --- | --- |
| **Less than every 5 years** | 0 | 0 | 0 |
| **Every 1-5 years** | 0 | 0 | 2 |
| **Annually** | 0 | 1 | 0 |
| **Multiple times per year** | 2 | 86 | 2 |
| **Daily** | 1 | 0 | 5 |

#### Physical habitat scores

Table S2: Unweighted habitat maintenance scores

| **Activity** | **Never** | **Less than every 5 years** | **Every 1-5 years** | **Yearly** | **Multiple times per year** |
| --- | --- | --- | --- | --- | --- |
| **Bank trimming** | 1 | 2 | 3 | 4 | 5 |
| **In-channel weed cutting** | 1 | 2 | 3 | 4 | 5 |
| **Desilting** | 1 | 2 | 3 | 4 | 5 |
| **Dredging** | 1 | 2 | NA | NA | NA |

Table S3: Weighted habitat maintenance scores

| **Activity** | **Never** | **Less than every 5 years** | **Every 1-5 years** | **Yearly** | **Multiple times per year** |
| --- | --- | --- | --- | --- | --- |
| **Bank trimming** | 1 | 2 | 3 | 4 | 5 |
| **In-channel weed cutting** | 1 | 3 | 4 | 5 | 6 |
| **Desilting** | 1 | 4 | 5 | 6 | 7 |
| **Dredging** | 1 | 8 | NA | NA | NA |

#### Community analysis

Fish assemblage composition was analysed using non-metric multidimensional scaling (NMDS) based on Jaccard dissimilarities calculated from presence–absence eDNA detections. Analyses were conducted at both the collapsed site level and the individual visit level. To avoid circularity, eel detections were excluded from community matrices prior to ordination, with eel presence used only as a grouping variable. Differences in assemblage composition associated with eel presence, sampling round, and their interaction were tested using permutational multivariate analysis of variance (PERMANOVA; adonis2). Homogeneity of multivariate dispersion was assessed using betadisper. Convex hull ellipses were used to visualise group distributions in ordination space.

### Results

#### Non-eel fish species

The marine species detected were European smelt (*Osmerus eperlanus*; two sites), European flounder (*Platichthys flesus*; one site) and sand goby (*Pomatoschistus minutus*; two sites), while gudgeon (*Gobio gobio*; 36 sites) and stone loach (*Barbatula barbatula*; 34 sites) were the most prevalent rheophilic species. Eighty-one pumped catchments were inhabited by only eurytopic and/or limnophilic species, i.e. no European eel, marine or rheophilic species were detected.

Spined loach occurrence differed significantly among connectivity categories (Fisher’s exact test: p = 0.000387). Occurrence was approximately four times more likely at sites with manual connectivity (odds ratio = 4.05, 95% CI = 1.88–8.97; p = 0.00010), while occurrence was significantly lower at sites with no connectivity (odds ratio = 0.29, 95% CI = 0.10–0.79; p = 0.0117) and natural connectivity (odds ratio = 0.38, 95% CI = 0.14–0.97; p = 0.0290).

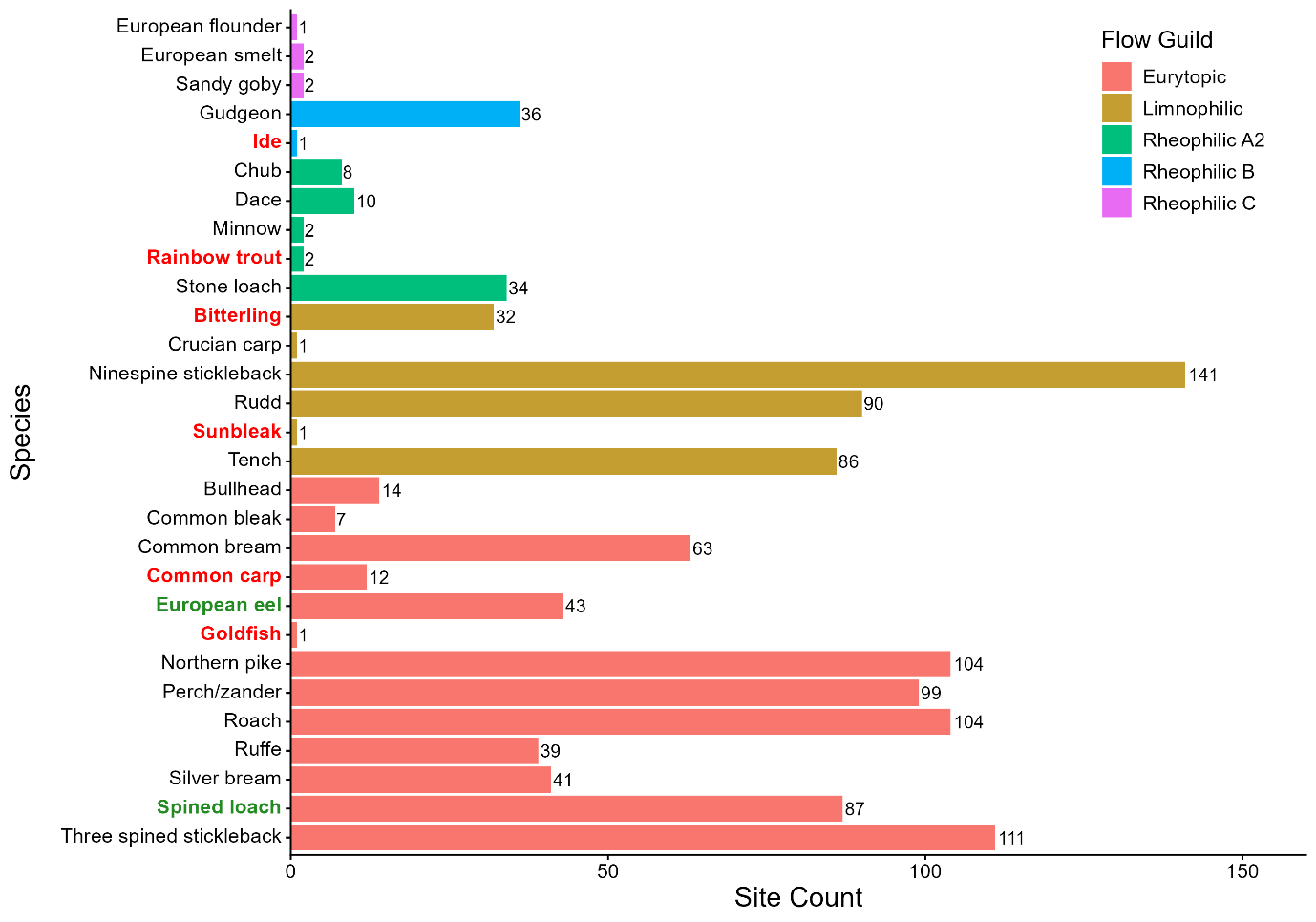

Figure S2. Numbers of sites fish species were detected at in the study regions, split by flow guilds. Red and green name labels denote non-native and conservation priority species, respectively.

#### Inter-sampling round effects on eel detection and overall fish richness

Eels were detected in 10/152 (6.6%), 10/107 (9.3%), 17/107 (15.9%) and 6/90 (6.7%) sites in the 1^st^, 2^nd^, 3^rd^ and 4^th^ sampling visits, respectively. There was limited visual separation of fish assemblages between eel-positive and eel-negative sites across sampling rounds, with substantial overlap in ordination space and ellipses largely encompassing both groups (Figure S3). Although PERMANOVA indicated a statistically significant effect of eel presence at the site level, the effect size was small (R² = 0.029, p = 0.007), indicating that eel presence explained only a small proportion of variation in community composition. Fish assemblages also differed significantly between sampling visits, but again with a small effect size (R² = 0.036, p = 0.001). A significant interaction between visit and eel presence was detected (R² = 0.057, p = 0.001), although this effect was also small, suggesting that the influence of eel presence on community composition varied across sampling rounds but remained weak overall. When analysed within individual visits, a significant eel-associated difference was observed only in visit 3 (R² = 0.049, p = 0.001), while other visits showed no significant effects.

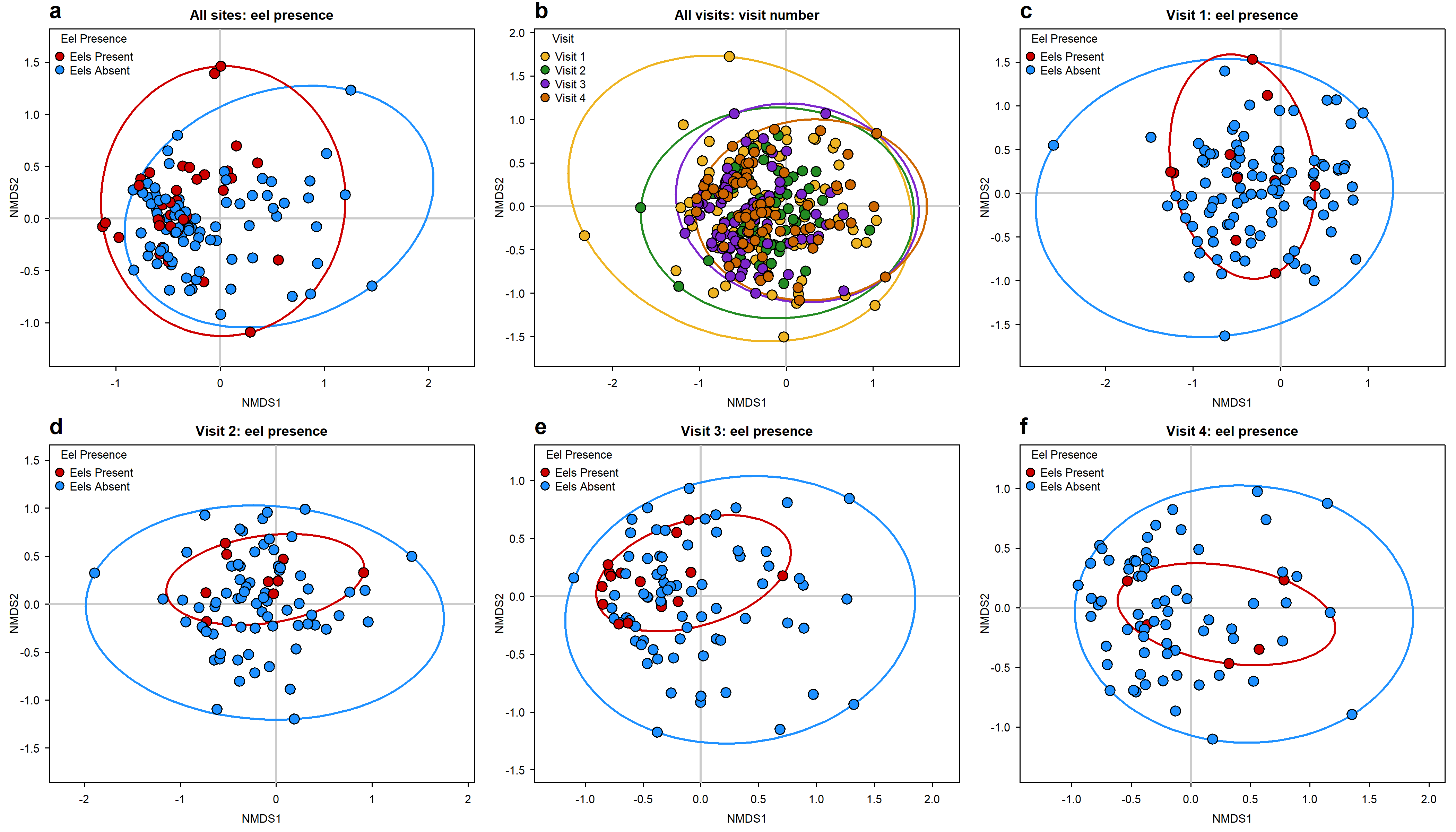

Figure S3. Non-metric multidimensional scaling (NMDS) ordination plots of fish assemblage composition based on presence–absence eDNA detections from fen sites. Panel (a) shows all sites grouped by eel presence or absence. Panel (b) shows all visit-level samples grouped by sampling visit. Panels (c–f) show R1, R2, R3 and R4, respectively, with sites grouped by eel presence or absence. Red points indicate eel-positive sites and blue points indicate eel-negative sites. Convex hull ellipses represent the extent of each group in ordination space.
